# Designed Endocytosis-Triggering Proteins mediate Targeted Degradation

**DOI:** 10.1101/2023.08.19.553321

**Authors:** Buwei Huang, Mohamad Abedi, Green Ahn, Brian Coventry, Isaac Sappington, Rong Wang, Thomas Schlichthaerle, Jason Z. Zhang, Yujia Wang, Inna Goreshnik, Ching Wen Chiu, Adam Chazin-Gray, Sidney Chan, Stacey Gerben, Analisa Murray, Shunzhi Wang, Jason O’Neill, Ronald Yeh, Ayesha Misquith, Anitra Wolf, Luke M. Tomasovic, Dan I Piraner, Maria J. Duran Gonzalez, Nathaniel R. Bennett, Preetham Venkatesh, Danny Satoe, Maggie Ahlrichs, Craig Dobbins, Wei Yang, Xinru Wang, Dionne Vafeados, Rubul Mout, Shirin Shivaei, Longxing Cao, Lauren Carter, Lance Stewart, Jamie B. Spangler, Gonçalo J.L. Bernardes, Kole T. Roybal, Per Greisen, Xiaochun Li, Carolyn Bertozzi, David Baker

**Affiliations:** Department of Biochemistry, University of Washington, Seattle, WA, USA; Institute for Protein Design, University of Washington, Seattle, WA, USA; Department of Bioengineering, University of Washington, Seattle, WA, USA; Department of Chemistry, Stanford University, Stanford, CA, USA; Howard Hughes Medical Institute, University of Washington, Seattle, WA, USA; Department of Molecular Genetics, University of Texas Southwestern Medical Center, Dallas, TX, USA; Novo Nordisk A/S, Måløv, Denmark; Departments of Biomedical Engineering and Chemical and Biomolecular Engineering, Johns Hopkins University, Baltimore, MD, USA; Medical Scientist Training Program, Johns Hopkins University School of Medicine, Baltimore, MD, USA; Department of Microbiology and Immunology, University of California San Francisco; Harvard Medical School, Harvard University, Boston, Massachusetts, USA; Division of Biology and Bioengineering, California Institute of Technology, Pasadena, CA, USA; School of Life Sciences, Westlake University, Hangzhou, China; Yusuf Hamied Department of Chemistry, University of Cambridge, Cambridge, UK; Instituto de Medicina Molecular João Lobo Antunes, Faculdade de Medicina, Universidade de Lisboa, Lisbon, Portugal; Howard Hughes Medical Institute, Stanford, CA, USA; Sarafan ChEM-H, Stanford University, Stanford, CA 94305, USA

## Abstract

Endocytosis and lysosomal trafficking of cell surface receptors can be triggered by interaction with endogenous ligands. Therapeutic approaches such as LYTAC^1,2^ and KineTAC^3^, have taken advantage of this to target specific proteins for degradation by fusing modified native ligands to target binding proteins. While powerful, these approaches can be limited by possible competition with the endogenous ligand(s), the requirement in some cases for chemical modification that limits genetic encodability and can complicate manufacturing, and more generally, there may not be natural ligands which stimulate endocytosis through a given receptor. Here we describe general protein design approaches for designing endocytosis triggering binding proteins (EndoTags) that overcome these challenges. We present EndoTags for the IGF-2R, ASGPR, Sortillin, and Transferrin receptors, and show that fusing these tags to proteins which bind to soluble or transmembrane protein leads to lysosomal trafficking and target degradation; as these receptors have different tissue distributions, the different EndoTags could enable targeting of degradation to different tissues. The modularity and genetic encodability of EndoTags enables AND gate control for higher specificity targeted degradation, and the localized secretion of degraders from engineered cells. The tunability and modularity of our genetically encodable EndoTags should contribute to deciphering the relationship between receptor engagement and cellular trafficking, and they have considerable therapeutic potential as targeted degradation inducers, signaling activators for endocytosis-dependent pathways, and cellular uptake inducers for targeted antibody drug and RNA conjugates.

## Main Text

The endocytosis of many cell surface receptors is triggered by binding of their endogenous ligands which can shift the conformational or oligomerization state of the receptor^4^ and induce receptor clustering and adaptor protein recruitment^5,6^. Native endocytosis inducing ligands have been utilized to enhance intracellular delivery and prompting extracellular protein degradation^1–3^. While powerful, these approaches have the limitations that native ligands can trigger off-targeting signaling^3,7^, their binding sites may be occupied by existing ligands, and instability and in some cases the need for modification can complicate manufacturing^8^. Bio-orthogonal inducers of endocytosis could have therapeutic utility not only for targeted degradation but also for initiating signaling through pathways involving endocytosis^4^, and provide powerful tools for investigating the association between cellular trafficking and receptor structures. Antibodies have been identified that stimulate endocytosis, but this can require considerable empirical screening for any target receptor^9,10^. To date there does not exist a completely synthetic and systematic approach to design such orthogonal and synthetic activators of cellular trafficking.

We reasoned that de novo protein design could enable the creation of bio-orthogonal endocytosis inducing proteins that avoid the above limitations by using strategies customized for the target receptor. For receptors such as sortillin and the transferrin receptor, that constitutively traffic between the cell surface and the endosome/ lysosome, binding to a site on the receptor non-overlapping with the native ligands could be sufficient (**Fig. 1a**). For receptors such as the IGF-2R receptor, for which conformational change triggers endocytosis, binding must induce rearrangement of receptor extracellular domains, while for others, such as ASGPR, where endocytosis is stimulated by clustering, binding should induce oligomerization. Fusion of designed proteins with these properties to a second target binding protein could promote endocytosis and lysosomal trafficking of the target. We set out to design such endocytosis targeting proteins, which we call EndoTags, for all four receptor systems, and to explore their utility for modulating protein degradation and cellular signaling.

**Figure 1:**
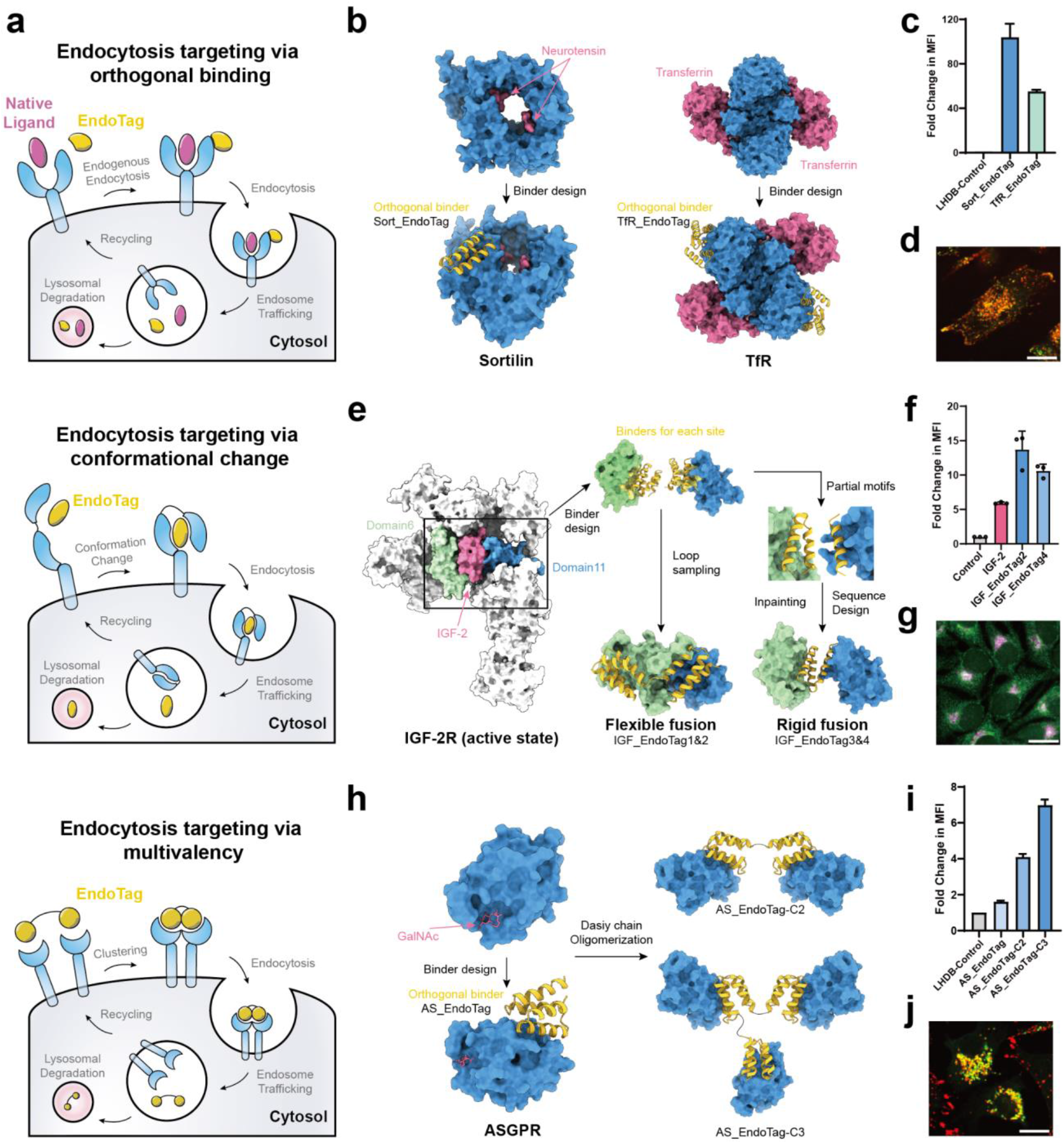
Design strategies for endocytosis-triggering EndoTags. **a,** Schema of designed endocytosis mechanisms. (top) Design of binding to constitutively cycling receptors at sites that do not overlap with those of natural ligands to avoid competition. (middle) Design of binders that trigger endocytosis by eliciting a conformational change in the receptor. The EndoTag binds at two distinct epitopes on the target and actively triggers the conformational change. (bottom) Designed endocytosis via receptor clustering. The multivalent EndoTag clusters multiple copies of the target receptor and induces endocytosis. **b,** Design strategy for Sortilin and TfR EndoTags. (left) Starting from the crystal structure of Sortilin^44^, minibinders were designed to bind at a site not overlapping with the native ligand neurotensin. (Right) A designed binder to TfR binds at a site distant from that of transferrin^45^. **c,** Cellular uptake of Sortilin and TfR EndoTags. U-251MG cells were treated with 100nM AF647-labelled Sortilin-EndoTags or TfR-EndoTags for 2h. After washing 3 times, cellular uptake was measured by flow cytometry. The data were normalized with respect to a control group treated with 100nM AF647-labelled LHDB (supposed no endocytosis). **d,** Confocal imaging of Sort_EndoTag (red) and a lysosomal marker (green). 500nM AF647 labelled Sort_EndoTag was incubated with U-251MG cells for 24h. The lysosomes were stained with AF488-labelled Lysotracker. **e,** Design strategy for IGF_EndoTags. Starting from the structure of IGF-2 in complex with IGF-2R, de novo minibinders were generated and screened against the IGF-2 binding sites at IGF-2R domain 6 and domain 11, separately. Individual binders for each domain (D6mb and D11mb) were fused with flexible linkers or a rigid fusion interdomain connection. For flexible fusion, multiple linker lengths and fusion directions were sampled. For rigid fusion, the two major binding helices from D6mb and one major binding helix from D11mb were extracted as starting motifs. With protein Inpainting, geometries and fusion orders were sampled, and ranked based on Rosetta and alphafold2 metrics. **f,** Cellular uptake of IGF_EndoTags. Jurkat cells were treated with biotinylated 100 nM IGF_EndoTags or IGF-2, and 33 nM Strapavidin-AF647 for 24h. After washing 3 times, cellular uptake was measured by flow cytometry. The data were normalized with the control group treated with 33 nM Strapavidin-AF647 alone. **g,** Microscopy imaging of IGF_EndoTag1 (pink) co-localization with lysosomes (green). Hela cells were incubated with 100nM biotinylated EndoTag1 and 33nM Strapavidin-AF647 for 2h. After washing and fixing the cells, the cells were stained with anti-LAMP2A antibody followed by fluorophore-labeled secondary antibody and DAPI. **h,** Design strategy for ASGPR EndoTags. Based on the crystal structure of ASGPR^46^, binders were designed and then connected with flexible linkers. **i,** Cellular uptake of ASGPR EndoTags. Hep3B cells were treated with 100 nM AF-647 conjugated ASGPR EndoTags for 24h. After washing 3 times, the cellular uptake was measured by flow cytometry. The data were normalized with respect to a control group treated with 100nM AF647-labelled LHDB (supposed no endocytosis). **j,** Confocal imaging of Sort_EndoTag (red) with lysosome (green). 500nM AF647 labelled Sort_EndoTag was incubated with Hep3B cells for 24h. The lysosomes were stained with AF488-labelled Lysotracker. For **d**,**g**,**j**, the scale bar indicates 20um. For **c,f,i**, the mean values were calculated from triplicates, and error bars represent standard deviations.

To enable tissue-specific control over endocytosis for downstream applications, we selected target receptors with distinct tissue expression profiles. IGF-2R is expressed in most tissues, ASGPR is expressed primarily in the liver, TfR and Sortilin are both abundant in the brain, where TfR has broader expression in liver and muscles while Sortilin distributes in the spinal cord^11,12^. We developed customized design approaches to optimize endocytosis through each of these receptor systems, as described in the following sections.

### Design of EndoTags orthogonal to endogenous ligands

Tfr and Sortilin constitutively cycle between the cell surface and intracellular compartments. Thus, for these receptors, the challenge is not to actively induce endocytosis, but to bind to the receptor at a site that does not compete for the natural ligand, which could have undesired side effects and reduced efficiency. De novo protein design has the advantage of being able to target binders to specific sites of interest on a target^13–15^, and thus is well suited to designing protein binders that target receptor sites that do not overlap with those of native ligands.

Sortilin is a rapid trafficking receptor with considerable expression in the neural system which plays a role in lysosomal targeting of neurotrophin^16,17^. We sought to design protein binders of Sortilin at binding sites not overlapping with native ligands including neurotrophin (**Fig. 1a,b**). We used Rosetta de novo binder design^13^ (**Extended Data** Fig. 1a) to generate 21,000 binders against the epitope (F92, V93, T546, T559, and T561) of Sortilin (**Extended Data** Fig. 1b-c), while avoiding any sites of known interactions and of significant structural change at low pH^16^. After experimental screening with yeast display and BLI (Biolayer Interferometry), the best candidate (Sortmb) bound Sortilin with 21nM affinity (**Extended Data** Fig. 2g). To validate the binding accuracy for Sortmb, we designed and expressed four Sortilin variants that contain N-linked glycan close to the designed Sortmb interface. In the yeast display assay, the binding affinity of Sortmb1 was reduced (**Extended Data** Fig. 8a-f), indicating that N-linked glycans are present in the designed interface and the computationally designed binder was binding to the intended interface. We next evaluated the ability of the binder (Sort_EndoTag) to drive internalization by flow cytometry. After incubation of 200nM fluorescence-labeled Sort_EndoTag with U-251 MG glioblastoma cells for two hours at 37°C followed by extensive washing, there was a ninety fold increase in fluorescence compared to fluorophore conjugated control (**Fig. 1c**). Confocal imaging indicated co-localization of the Sort_EndoTag with a lysosomal marker after 24 hours incubation in U-251MG cells (**Fig. 1d, Extended Data** Fig. 5a).

We applied a similar orthogonal binding strategy with TfR (**Fig. 1b**) whose native function is to transport iron-bound transferrin into cells and across the blood brain barrier^12,18^. We previously reported the design of TfR binders with high binding affinity at a binding site distant from the transferrin binding site^19^; here we explore the use of this binder as an EndoTag (TfR_EndoTag). We found that the TfR_EndoTag directed endocytosis in U-251MG glioblastoma cells, with a fifty fold increase in cellular uptake over control after two hour incubation (**Fig. 1c**). Confocal imaging again indicated lysosome targeting of the TfR_EndoTag after 24 hours incubation in U-251MG cells (**Extended Data** Fig. 5b).

Taken together, these results with Sortilin and Tfr indicate that binding to bio-orthogonal sites with single high-affinity binding domains is an effective strategy for hijacking constitutively endocytosing receptors. Our bio-orthogonal binder design approach could be readily generalized to a wide range of other tissue specific targets.

### Design of EndoTags for triggering conformational change

IGF-2R is a large cell surface receptor that rapidly transports its natural ligands IGF-2 and M6P to the lysosome for degradation^1^. IGF-2R has high abundance across tissues, and considerable overall lysosomal trafficking capacity which has made it an attractive target for protein degradation approaches. Structure investigation suggests that IGF-2 binding induces a conformational change in IGF-2R which brings together domain 6 (D6) and domain 11 (D11), and promotes dimerization of the receptor^20^. With the ability to design de novo binders at arbitrary interfaces^13^, we hypothesized that a designed binding protein that brings together domain 6 and 11 could similarly trigger IGF-2R endocytosis and lysosomal targeting without triggering off-target signaling activation like IGF-2^7^.

We sought to use the Rosetta RIFdock method^13,14^ to design small proteins that bind to D11 and D6 of IGF-2R independently. Inverse rotamer fields were generated around residues interacting with IGF-2R in both domains (**Extended Data** Fig. 1d-g), and scaffolds were docked into these fields with RIFdock^21^. After sequence optimization and refinement with RosettaFastdesign, the best scoring binding motifs were identified and grafted back on to the scaffold library with RosettaMotifgraft^15^, followed by another round of RosettaFastdesign. Genes encoding the top-ranked 15,000 designs based on Rosetta interface metrics (ddg, contact_molecular_surface)^13^, were synthesized on an oligonucleotide array, transformed into a yeast display vector, and fluorescence-activated cell sorting (FACS) was carried out using biotinylated human IGF-2R domain 6 and domain 11. After next-generation sequencing, we identified one high-affinity candidate binder for domain 6 and two candidate binders for domain 11; these were expressed and purified in E coli. The IGF-2R domain 6 binder (D6mb) affinity of 41nM KD was measured using biolayer interferometry (BLI) (**Extended Data** Fig. 2a), and the best domain 11 minibinder (D11mb) bound IGF-2R domain 11 at an affinity of 190nM KD (**Extended Data** Fig. 2b). After affinity optimization with mutation library screening, the highest affinity variant for D11 (D11mb2) bound domain 11 at 6.5nM KD in BLI (**Extended Data** Fig. 2c).

We next sought to develop IGF-2R EndoTags (IGF_EndoTags) utilizing the distinct IGF-2R D6 and D11 minibinders (**Fig. 1e**). Initially, we explored the use of flexible fusions between D11mb and D6mb, with different loop lengths and domain orders. We then expressed these fusions in E. coli and incorporated an avi-tag for biotinylation. Finally, we evaluated their cellular uptake in Jurkat cells by utilizing Alexa-647 (AF-647) and flow cytometry. A construct connecting D11mb and D6mb with a GGS linker (D11mb-GGS-D6mb, IGF_EndoTag1) was the most readily uptaken, with increased cell associated fluorescence signal over native IGF-2 or D6mb/D11mb alone in Jurkat cells (**Extended Data** Fig. 4c). Co-localization imaging indicated that IGF_EndoTag1 is targeted to lysosomes (**Fig. 1g**): this synthetic ligand thus recapitulates the trafficking of the endogenous IGF-2 ligands. Longer linkers decreased the uptake level, while a shorter GS linker abolished all uptake (**Extended Data** Fig. 4b), suggesting the orientation and distance of the two binding domains modulates the IGF-2R endocytosis process. Constructs with two copies of one mini-binder (D6mb-linker-D6mb, D11mb-linker-D11mb did not display any cellular uptake (**Extended Data** Fig. 4a); engagement of both domains (likely driving their reorientation within the receptor structure) appears to be necessary to trigger efficient cellular uptake. Substitution of D11mb with the higher affinity variant D11mb2 in IGF_EndoTag1 (generating IGF_EndoTag2) increased internalization to 2 fold compared to native IGF-2 in Jurkat cells (**Fig 1f**); IGF_EndoTag2 was clearly detectable in lysosomes after a 30 minute incubation, unlike IGF-2 or IGF_EndoTag1 (**Extended Data** Fig. 4d-e).

We reasoned that still more potent stimulation of endocytosis could be achieved by two domain constructs in which the individual domains are rigidly fused to each other to drive specific receptor conformational changes. We generated such fusions by using RFInpainting^22^ to generate three-helix bundles combining the major interface helix from D11mb and two interface helices from D6mb (**Fig. 1e**, **Extended Data** Fig. 3). Designed residues within 3 angstroms (A) of the receptor were kept fixed, the D11mb helix was randomly perturbed by rigid body translations (up to 5 angstroms) and rotations (up to 10 degrees), the two chains were connected by inpainting, and the sequence of the fusion designed in the context of IGF-2R using ProteinMPNN^23^ keeping residues interacting with the receptor constant. Models with RosettaFold LDDT metrics (>0.5) for the inpainted region and Alphafold2^24^ (AF2) structure predictions for the fusions matching the design models (RMSD < 2.0) were experimentally screened for binding with yeast display against both IGF-2R domains. Out of 170 screened designs, the 8 most enriched candidates were selected for protein expression and BLI affinity evaluation. Different designs had distinct affinities for domains 6 and 11; for example, IGF_EndoTag3 possessed strong binding affinity to both domains (6nM for domain 6 and 190nM for domain 11, **Extended Data** Fig. 2d), while EndoTag4 bound more tightly to domain 6 (15nM for domain 6 and 4.3µM for domain 11, **Extended Data** Fig. 2e). In cellular uptake assays, IGF_EndoTag3 had internalization activity equivalent to IGF-2, and IGF_EndoTag4 showed 2 fold higher internalization than IGF-2 (**Extended Data** Fig. 4c) and co-localized with lysosomes within 30 minutes as we observed in co-localization imaging (**Extended Data** Fig. 4f).

### Design of EndoTags for inducing receptor clustering

The endocytosis of receptors such as ASGPR and EGFR is stimulated through dimerization or oligomerization^25^. ASGPR is a liver-specific ligand shuttling receptor to transport GalNAc labeled proteins into lysosomes for clearance^26^. Multivalent GalNAc ligands have been used for multiple liver-specific degradation applications^2,27^ and RNA delivery platforms^28^. However, these require chemical modification and hence are not genetically encodable, and must compete with native ligands which raises challenges.

We designed binders to ASGPR that do not overlap with the glycan binding sites (**Extended Data** Fig. 1h**&i**) using an augmented version of the Rosetta design approach described above that uses ProteinMPNN^23^ for sequence design and Alphafold2^18^ for design evaluation. 2689 designs passed Alphafold2 pae_interaction cutoff and were selected for experimental screening. After yeast display enrichment and NGS, we identified 4 candidates that bound to ASGPR on the yeast surface. Following expression and purification from E coli, biolayer interferometry showed that the highest affinity of these, (ASmb1), binds to ASGPR with an affinity of 2.7 µM (**Extended Data** Fig. 2f).

To stimulate ASGPR endocytosis through clustering, we connected two or three ASmb1 domains with GS linkers to generate ASGPR EndoTags, which we refer to as AS_EndoTags-2C and AS_EndoTags-3C (**Fig. 1h**). Cellular uptake assays in Hep3B cells indicated that multivalent AS_EndoTags-2C and AS_EndoTags-3C were endocytosed more efficiently than monomeric ASmb1 (following a two hour incubation and extensive washing, 2.5-fold and 4.5-fold more fluorescence was associated with cells, respectively (**Fig. 1i**)). Confocal imaging showed that AS_EndoTag-3C strongly co-localized with lysosomes after 24h (**Fig. 1j**, **Extended Data** Fig. 5c).

### Cell surface receptor degradation

Targeted degradation of receptors and extracellular soluble proteins is a promising therapeutic strategy for cancer, autoimmune diseases and neurodegenerative diseases. Lysosome-targeting Chimeras (LYTACs) utilize mannose-6-phosphonate (M6Pn) ligands which triggers lysosomal delivery and degradation of the targeted proteins through the IGF-2R^1,29^, or N-acetylgalactosamine (GalNAc) to trigger the ASGPR lysosomal trafficking pathway^2^. While very promising, the LYTAC approach is hindered by the reliance on existing native ligands and by the sophisticated chemistry required to generate multivalent modifications that increase endocytosis potency, complicating their manufacturing. Given their potent and rapid endocytosis and lysosomal targeting ability, we hypothesized that the fusion of EndoTags with POI-specific binders to generate protein-LYTACs (pLYTAC) (**Fig. 2a**), could provide an orthogonal and genetically-encoded approach for efficient extracellular protein degradation, and the different tissue distributions of the different receptors could allow targeting of degradation to different tissues.

**Figure 2.**
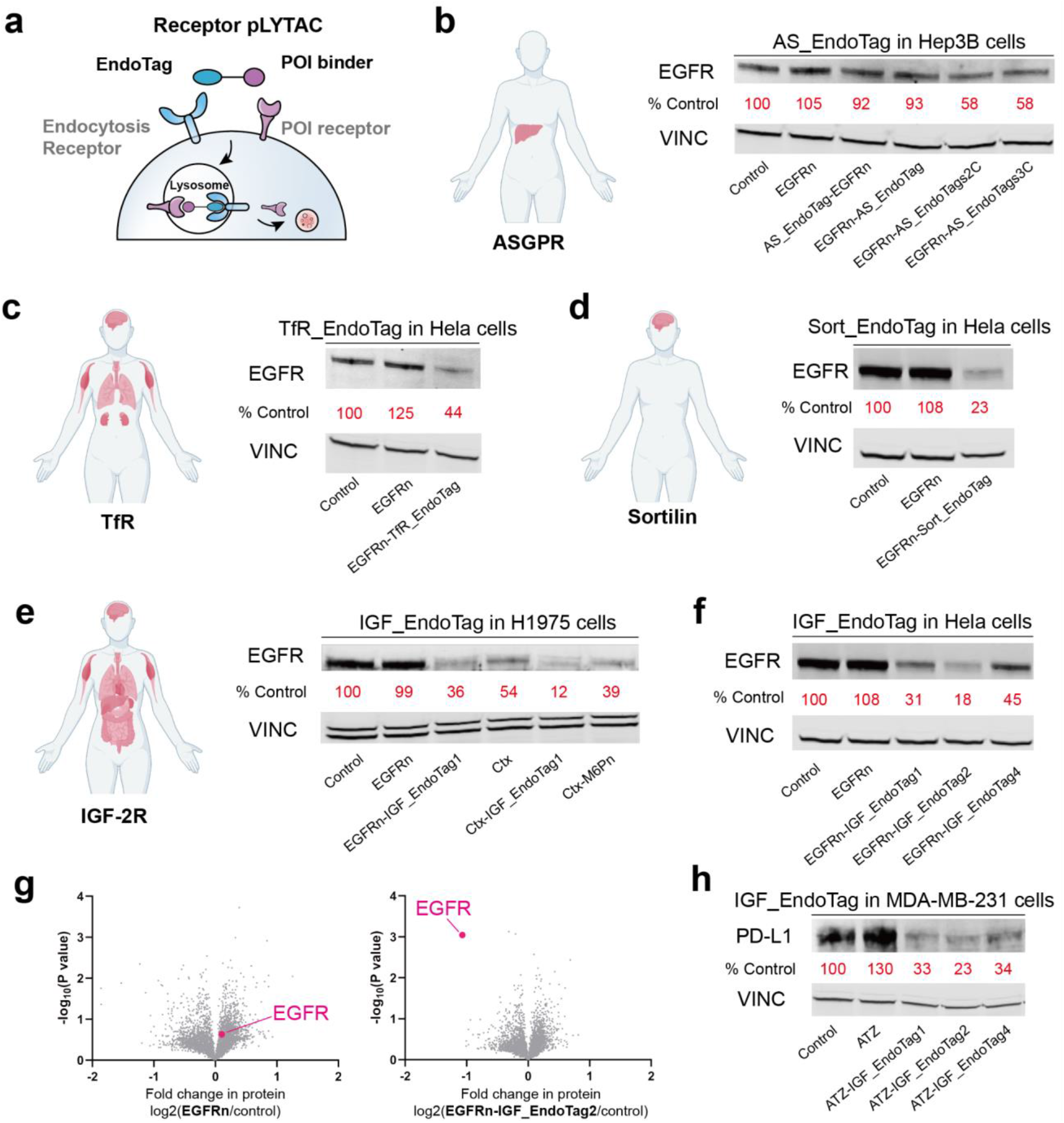
Surface receptor degradation with tissue-specific pLYTACs. **a,** Schema of tissue-specific pLYTACs for receptor degradation. **b,** ASGPR. Left: Tissue expression of ASGPR. Right: Western blot analysis of total EGFR levels in Hep3B cells after treatment with 200 nM EGFRn or EGFRn-AS_EndoTag for 48 h. **c,** TfR. Left: Tissue expression of TfR. Right: Western blot analysis of total EGFR levels in Hela cells after treatment with 200 nM EGFRn or EGFRn-TfR_EndoTag for 48 h. **d,** Sortilin. Left: Tissue expression of Sortilin. Right: Western blot analysis of total EGFR levels in Hela cells after treatment with 200 nM EGFRn or EGFRn-Sort_EndoTag for 48 h. **e,** IGF-2R. Left: Tissue expression of IGF-2R. Right: Western blot analysis of total EGFR levels in H1975 cells after treatment with 200 nM of EGFRn/CTX with/without the fusion with EndoTag1 or M6Pn for 48 h. **f,** Western blot analysis of total EGFR levels in Hela cells after treatment with 200 nM EGFRn or EGFRn-IGF_EndoTags for 48 h. **g,** Quantitative proteomics analysis of protein abundance in H1975 cells after treatment with EGFRn or EGFRn-IGF_EndoTag2 for 48h relative to untreated cells. Data are the mean of three biological replicates. **h,** Western blot analysis of total PD-L1 levels in MDA-MB-231 cells after treatment with 200 nM of ATZ or ATZ-pLYTACs for 48 h.

To explore the potential of pLYTACs in promoting protein degradation, we investigated their ability to target and degrade cell surface receptors. We started by focusing on the Epidermal Growth Factor Receptor (EGFR), which is often overexpressed in various cancer types and plays an important role in regulating cell proliferation^30^. We first assessed the degradation efficiency of the liver specific AS_EndoTags. Consistent with the cellular uptake results (**Fig. 1k**), introduction of EGFRn fusions to AS_EndoTags-2C and AS_EndoTags-3C resulted in a 40% decrease in total EGFR levels, while fusions to the monomeric ASGPR binder had little effect (**Fig. 2b**). Thus, designed protein mediated clustering of ASGPR can drive lysosomal targeting and cargo degradation, and the ASGPR-EndoTags could serve as liver-specific targeted degraders. To generate EGFR-pLYTACs targeting in the brain, we fused TfR_EndoTag and Sort_EndoTag with a minibinder targeting EGFR N-terminal (EGFRn)^13^. We observed efficient clearance of EGFR in HeLa cancer cells after 48h incubation, with 55% reduction of EGFR by EGFRn-TfR_EndoTag (**Fig. 2c**) and 78% reduction of EGFR by EGFRn-Sort_EndoTag western blot (**Fig. 2d**). Given the abundant expression of the corresponding receptors in the brain, both TfR_EndoTag and Sortilin_EndoTag could function as pLYTACs for neurodegenerative disease applications.

We next sought to make systemically active pLYTACs that act through the ubiquitously expressed IGF-2 receptor. To this end, we fused IGF_EndoTags to EGFRn, and found that in H1975 and HeLa cells, EGFR was effectively cleared from the whole cell lysate, as evidenced by western blot analysis (**Fig. 2e&f, Extended Data** Fig. 6b). EGFRn-IGF_EndoTag2 was the most effective, leading to more than 80% clearance of EGFR, while EGFRn-IGF_EndoTag1 resulted in 70% clearance. The flexible EGFRn-IGF_EndoTag1&2 had higher degradation capacity than the rigid constructs (IGF_EndoTag3&4) (**Fig. 2f**), perhaps due to better interactions with EGFR. Mass spectrometry-based proteomic analyses confirmed that EGFRn-IGF_EndoTag2 and EGFRn-IGF_EndoTag1 significantly reduced EGFR levels in both Hela and H1975 cells without affecting IGF-2R levels (**Fig. 2g, Extended Data** Fig. 6e); EGFRn without EndoTag had no effect on EGFR levels (**Fig. 2g, Extended Data** Fig. 6d).

To compare with the original M6P-based LYTACs, we generated genetic fusions of EndoTag1 with Cetuximab (CTX), a clinically-approved therapeutic antibody that targets EGFR with high affinity^31^. In H1975 cells, CTX-IGF_Endotag1 led to more effective degradation of EGFR than the M6P-based LYTAC (**Fig. 2e**)^1^. Proteomic analyses demonstrated that CTX-IGF_Endotag1 elicited a significantly greater reduction in EGFR levels compared to CTX alone (**Extended Data** Fig. 6f-g), with little effect on IGF-2R levels. CTX has a higher affinity for EGFR, and hence required a lower dose for effective degradation: CTX-IGF_Endotag1 achieved 85% degradation at a concentration of 10nM (**Extended Data** Fig. 6a).

We next investigated targeted degradation of Programmed Death-Ligand 1 (PD-L1), an immune checkpoint in cancer immunotherapy, and therefore an attractive target for this approach^32^. We genetically fused pLYTACs at the C-term of Atezolizumab (Atz), a potent PD-L1 inhibitor^33^, then tested PD-L1 clearance ability. All of the Atz-EndoTags effectively degraded PD-L1 on the cell surface of MDA-MB-231 cells within 4 hours, with EndoTag2 achieving the highest clearance percentage of 90% after 24 hours of incubation (**Extended Data** Fig. 6h). Western blot analysis of the whole cell lysis indicated that Atz-EndoTag2 molecules were able to deplete 77% of PD-L1 in the whole cell after 48h incubation (**Fig. 2h**). Like PD-L1, cytotoxic T lymphocyte antigen 4 (CTLA4) is an immune checkpoint component for which inhibitors have shown promising antitumor effects^18,19^. To target CTLA4, we generated a genetic fusion between EndoTag1 and a minibinder against CTLA4 (CTLA4mb) [permission of unpublished] which resulted in a 45% decrease of CTLA4 in Jurkat-CTLA4 cells after only 3 hours (**Extended Data** Fig. 6c).

### Clearance of soluble proteins

We next investigated the ability of EndoTags to degrade targeted soluble proteins as soluble pLYTACs (**Fig. 3a**). As a first proof of concept we employed the nanomolar affinity de novo designed protein heterodimer LHDA and LHDB, which has been used as the basis for synthetic signaling systems^36^, [accompanying manuscript]). We fused LHDA to IGF_EndoTags, and LHDB to AF647, and found that the EndoTags significantly enhanced the uptake of LHDB-AF647 in both Jurkat and K562 cells (**Fig. 3b, Extended Data** Fig. 7a**&b**), with IGF_EndoTag3 producing a remarkable 40-fold increase in mean fluorescence intensity (MFI) compared to the control. Incubation of Jurkat cells with 100nM LHDA-IGF_EndoTag3 resulted in a 50% clearance of 100nM LHDB from the solution after 48h incubation in Jurkat cells (**Fig. 3c**; clearance may be limited by the number of available receptors).

**Figure 3:**
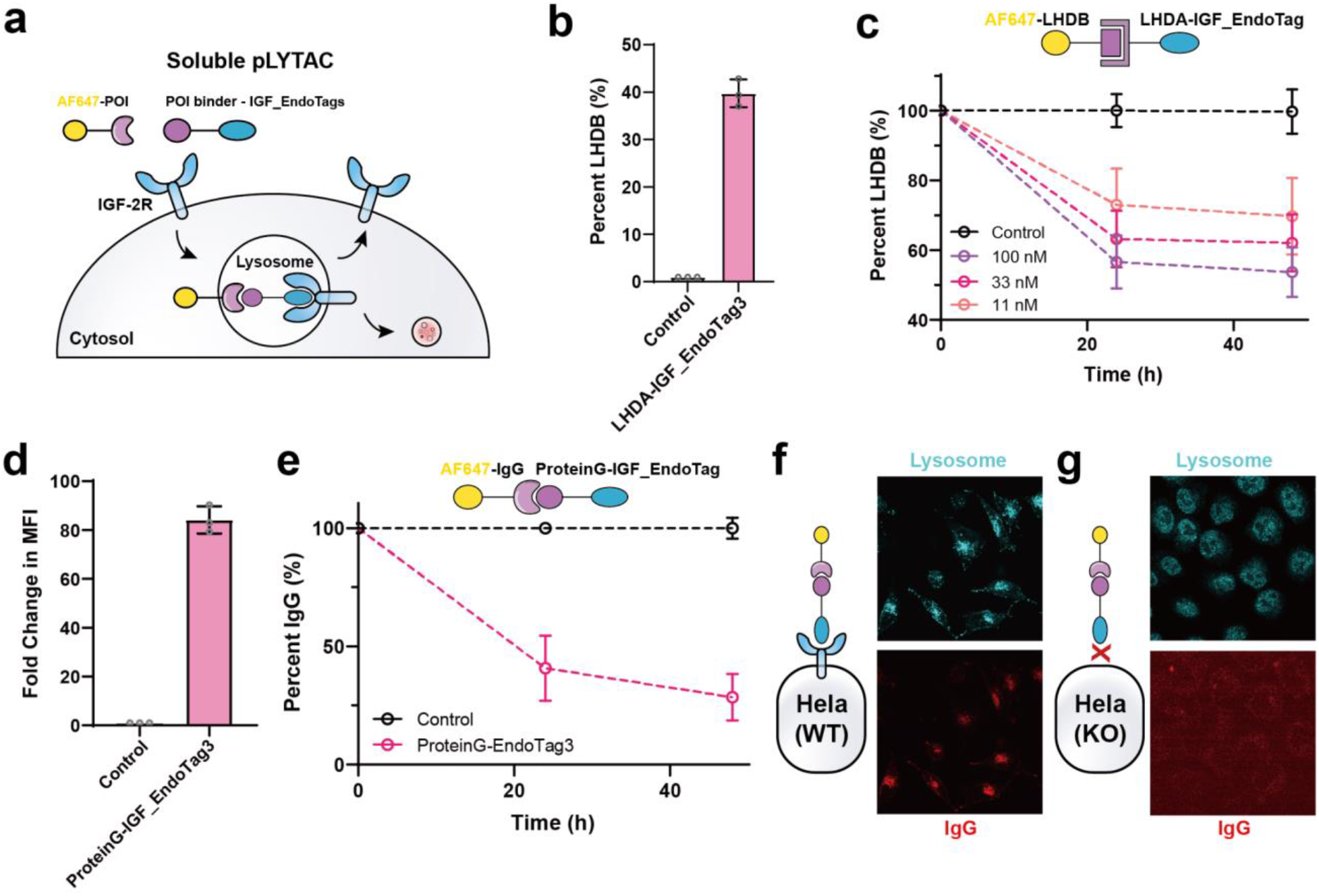
Clearance of soluble proteins by IGF-2R pLYTACs. **a**, Schema of soluble pLYTACs with IGF_EndoTags. **b**, Cellular uptake of LHDB-AF647 via LHDA-IGF_EndoTags in Jurkat cells. Cells were incubated with 33nM LHDB-AF647 with/wihtout 1uM LHDA-IGF_EndoTags for 24h, washed twice with cold PBS and analyzed by flow cytometry. **c**, Remaining supernatant LHDB-AF647 levels in Jurkat cells. Jurkat cells were incubated with 100nM LHDB-AF647 with/wihtout 500nM LHDA-IGF_EndoTags. At timepoints 24h, 48h, the cells were pelleted down, and supernatant IgG levels were quantified by Neo2 plate reader. The percentage of IgG level was normalized with the IgG alone control group. **d**, Cellular uptake of IgG-AF647 via proteinG-IGF_EndoTags in K562 cells. Cells were incubated with 33nM IgG-AF647 with/without 1uM proteinG-IGF_EndoTag3 for 24h, washed twice with cold PBS and analyzed by flow cytometry. The fold change in MFI (mean fluorescence intensity) was calculated by normalizing the IgG-AF647 alone group. **e**, Remaining supernatant IgG-AF647 levels in Jurkat cells. Jurkat cells were incubated with 133nM IgG-AF647 with/wihtout 100nM proteinG-IGF_EndoTag3. At timepoints 24h, 48h, the cells were pelleted down, and supernatant IgG-AF647 levels were quantified by Neo2 plate reader. The percentage of IgG-AF647 level was normalized with the IgG-AF647 alone control group at each time point. **f**, Confocal imaging of lysosome co-localization of IgG-AF647 with lysosome in Hela cells. **g**, Confocal imaging of lysosome co-localization of IgG-AF647 with lysosome in Hela (IGF-2R KO) cells. For **f**,**g**, the cells were incubated with 200nM IgG-AF647 and 1uM of proteinG-IGF_EndoTag3 for 24h, washed and stained with Lysotracker.

We next investigated the potential of the pLYTAC system to eliminate disease-relevant soluble protein targets such as autoantibodies that recognize self-antigens which have been linked to multiple autoimmune diseases^37^. We first tested whether the fusion of EndoTags with IgG binding protein G^38^ could clear IgG in solution. proteinG-IGF_EndoTags were generated by genetically fusing proteinG with various EndoTags and purified after expression in E coli. As a benchmark comparison to glycan-based LYTAC, ProteinG-M6Pn was generated by conjugating proteinG with azido-NHS ester followed by M6Pn-BCN peptide. All fusion proteins including EndoTag constructs together with proteinG-M6Pn were then incubated with IgG-AF647 and Jurkat cells. All candidates triggered considerable uptake of IgG in both Jurkat and K562 cells (**Extended Data** Fig. 7c**&e**). ProteinG-IGF_EndoTag3 elicited 2 fold higher cellular uptake of IgG than proteinG-M6Pn, leading to an overall 80 fold increase in IgG in K562 cells and 360 fold increase in Jurkat cells (**Extended Data** Fig. 7e). To quantify the clearance of IgG in the solution, we measured the fluorescence intensity in the cell culture supernatant normalized by that in control cells treated with IgG-AF647 alone. Incubation of 100nM proteinG-IGF_EndoTag3 with 133nM IgG resulted in depletion of 70% of the IgG after 48 hours in Jurkat cells (**Fig. 3e**). IGF_EndoTag3 elicited the higher clearance of IgG in Jurkat cells compared to the flexible IGF_EndoTag1&2, and a proteinG-M6Pn control (**Extended Data** Fig. 7d**&f**). IGF_EndoTag4 was less potent (**Extended Data** Fig. 7h). Utilizing confocal microscopy in HeLa cells, we observed enhanced co-localization of IgG with lysosomes following treatment with proteinG-IGF_EndoTag3 for 24 hours, indicating that the EndoTags actively facilitated the transportation of IgG to the lysosomes for degradation (**Fig. 3f**); this co-localization was not observed in HeLa cells not expressing IGF-2R (**Fig. 3g**).

### Logic gated targeted degradation and locally secretable degraders

Many targets of interest are expressed not only in diseased tissues but also on healthy cells, and the ability to target protein degradation to specific cell subpopulations could reduce toxicity. For example, in the case of cancer, target degradation conditioned on the presence of a specific marker in the tumor microenvironment could help avoid undesired effects on healthy cells. Such logic gated targeted degradation has not been achieved with current extracellular protein degradation systems. To address this limitation, we utilized the colocalization-dependent protein switches (Co-LOCKR) system, which functions as an AND logic gate only exposing a recruitment motif when two target cell markers are present on the same cell (**Fig. 4a**). We utilized Co-LOCKR to selectively degrade EGFR when HER2 was also present on the surface of cancer cells^39^. We fused EndoTag with Bcl2, which binds the Bim peptide exposed upon coincident binding in this version of Co-LOCKR, and evaluated EGFR degradation level in both HER2+ and HER2-cells. In K562 cells overexpressing both EGFR and HER2, Bcl2-IGF_EndoTag2 resulted in 80% degradation of EGFR, while in K562 cells expressing only EGFR without HER2, the EGFR level remained unchanged (**Fig. 4b**). Thus the EndoTag system can be precisely targeted to specific cells based on combinations of surface markers.

**Figure 4.**
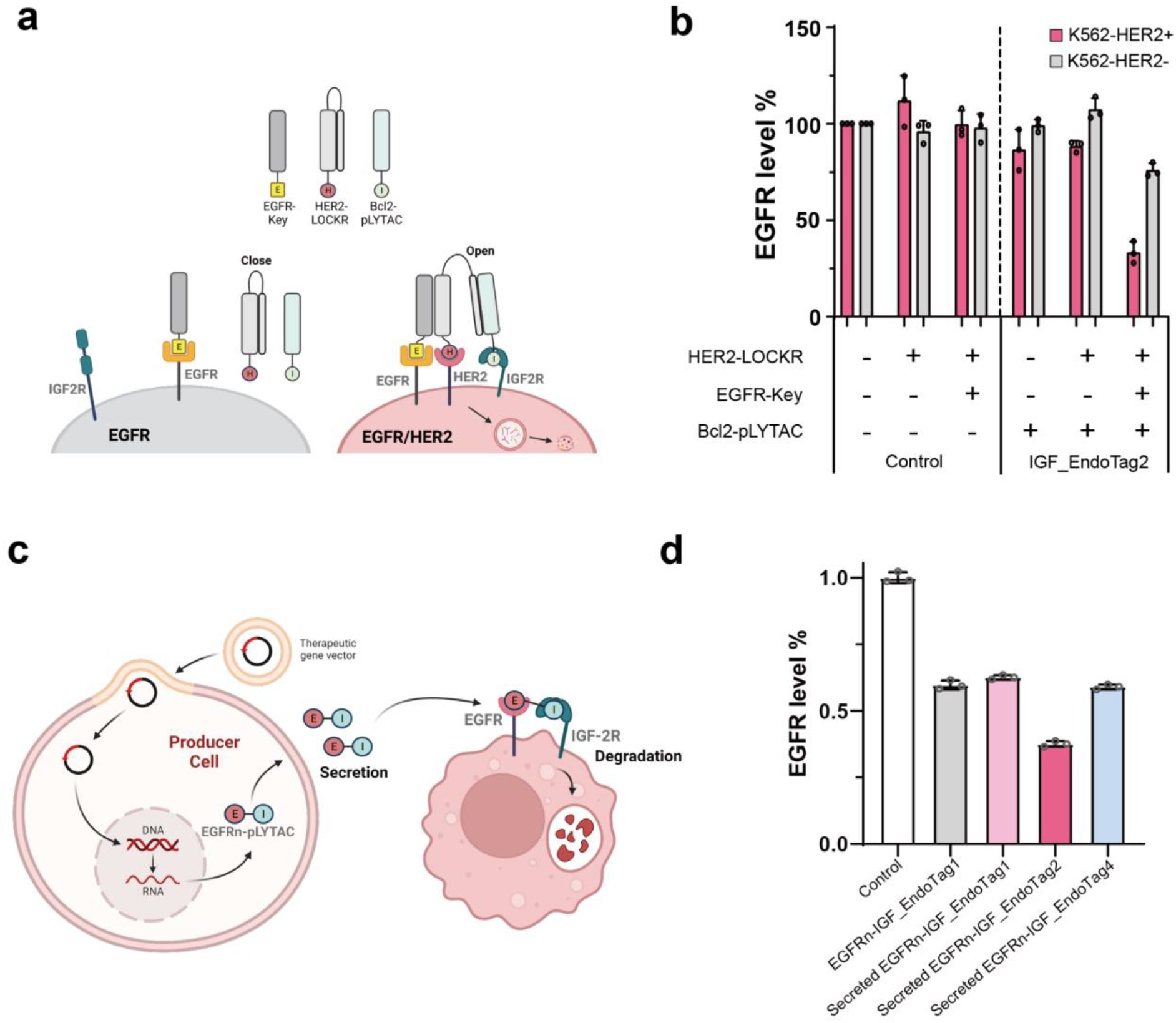
Logic gated targeted degradation and locally secretable degraders. **a,** Schema of AND-gate logic for EGFR degradation in presence of HER2**. b,** Flow cytometry quantification of EGFR level in cell surface. The K562-EGFR or K562-EGFR/HER2 cells were incubated with combinations of 100nM of EGFRn-Key, HER2-LOCKR, and Bcl2-EndoTag2 for 24h. **c,** Schema of secreted EGFR-pLYTAC. **d,** Flow cytometry quantification of EGFR level in cell surface treated with cell supernatant or exogenous EGFRn-IGF_EndoTag1 for 24h. For the secretion groups, Hela (IGF-2R KO) cells were transfected with viral vectors encoding the EGFRn-IGF_EndoTag1 or LHDA-pLYTACs. The supernatant of the cells were collected and incubated with K562-EGFR cells.

Another major challenge is the degradation of the degrader itself upon binding cells that express the target endocytosis receptor, which could increase the dose of drug required. In the context of adoptive cell therapies or mRNA delivery, the secretion of a degrader locally in response to stimuli could potentially overcome this challenge. Our system offers a solution as it is fully protein based and consequently can be locally secreted with high specificity and efficiency^40,41^. To investigate this possibility, we transiently transfected IGF-2R KO HeLa cells with plasmids encoding EGFRn-IGF_EndoTag, collected the cell supernatants, and incubated them with EGFR+ K562 cells while monitoring EGFR expression levels (**Fig. 4c**). Secreted EGFRn-IGF_EndoTag1 led to similar EGFR clearance as the exogenously added EGFRn-IGF_EndoTag1 (**Fig. 4d**), while secreted EGFRn-IGF_EndoTag2 was more potent, consistent with the results obtained with the purified proteins. Thus EndoTags remain functional when secreted from cells, providing a potential avenue for their use in adoptive cell therapy control systems.

## Conclusions

Targeted degradation has shown considerable promise as a therapeutic approach^1–3^ by using native ligands or chemical modifications to engage with rapidly trafficking cell surface receptors. Our designed EndoTag approach extends the power of this therapeutic modality in several ways. First, while native ligands can trigger off-target signaling and competition with endogenous proteins can reduce potency [permission of unpublished], our designed EndoTags, as illustrated by the Sortillin, TfR and ASGPR cases (**Fig. 1b&e&h**), can be targeted to sites on endocytosing receptors that are not bound by native ligands. Second, high valency chemical modification^1,2,42^ has been used to enhance endocytosis, but this complicates manufacturing^1,23,14^; as illustrated by our multi domain ASGPR EndoTags, small synthetic domains can be readily combined to create all protein receptor clustering and endocytosis stimulating proteins. The all-protein nature of our pLYTACs not only simplifies manufacturing, but also enables employment of targeted degradation approaches in adoptive cell therapies through secretion from engineered cells (**Fig. 4c**). The small, stable and readily producible synthetic ligands that could be useful not only for therapeutic applications but as molecular tools for probing how receptor conformations modulate cellular trafficking. Moving forward, there are many avenues to pursue using our computational design approach to generating EndoTag stimulated enhancers of cell surface receptor endocytosis and trafficking. First, there are likely many more receptor targets which can undergo rapid endocytosis upon suitable triggering at the cell surface; our ability to design endocytosis stimulators without requiring native ligands or identification of chemical modifications should enable utilization of the full range of these receptors to achieve more tissue restricted targeting and modulatable intracellular trafficking (different receptors likely will have different intracellular compartment residence times and transition dynamics). In addition to the targeted degradation application pursued here, such designed endocytosis stimulators could be of great utility for enhancing uptake of nucleic acid (siRNA for example) and small molecule drug conjugates. Second, as illustrated by the use of the LOCKR system to make logic gated protein degradation systems and the secretion of EndoTag-degrader constructs from cells, the robustness and modularity of de novo designed protein and capability for logic gated activation and cell based expression open the door to a wide range of more precise and controllable strategies for protein and cell based therapies.

## Methods

### Computational design of Sortilin minibinders

Using Rosetta-based binder design protocol^13^, 21,000 binders were generated to each of two sites on the Sortilin. The epitope of Site1 is comprised of five amino acids (Uniprot numbering: F92, V93, T546, T559, and T561) was chosen since it provided a modest patch of exposed hydrophobicity while avoiding any sites of known interactions. The selected epitope has the added feature that a binder to this location would be pH dependent since this region undergoes significant structural change at low pH^16^. As previously described^13^, we used a set of scaffold libraries to generate several million docks to each of the sites. As in the protocol, 100,000 were sub-selected and sequence designed. Helical motifs were extracted, and 3,000 designs were selected, grafted, and subjected to further design. Designs were filtered based on their Rosetta ddG and ContactMolecularSurface to the hydrophobic residues listed above. This resulted in 42,000 designs that were tested experimentally.

### Computational design of IGF-2R and ASGPR minibinders

The minibinders against IGF-2R domain 6 and 11 were computationally designed via a Rosetta-based approach previously described^13^. Briefly, the structures of IGF-2R domain 6 (PDB 6UM2) and IGF-2R domain 11 (PDB 1GP0) were refined using Rosetta Fastrelax with coordinate constraints. The residues at IGF-2 binding site for each domain were selected as “hotspot” residues. Helical protein scaffolds were docked against the hotspot residues via Patchdock followed by Rifdock protocol. After sequence optimization with Rosetta FastDesign and filtering with Rosetta interface metrics including ddg and contact_molecular_surface, the top candidates were then resamplered with Rosetta Motifgraft and FastDesign. Candidates passing previous filters were then filtered again with exposed hydrophobicity (sap_score) and optimized with a net-charge of -7.

The minibinders against ASGPR were designed with a Rosetta-based approach integrated with ProteinMPNN and Alphafold2. The crystal structure of ASGPR (PDB 5JQ1) was refined and helical protein scaffolds were docked against the exposed hydrophobic residues via Patchdock followed by Rifdock. The sequences were optimized with protein-MPNN and interface scores were calculated with Rosetta Fastrelax. The models were then predicted by Alphafold2 and scored after Fastrelax. Designs with pae_interaction<10 and relaxed_ddg<-40 were selected for resampling with another round of protein-MPNN prediction followed by Rosetta Fastrelax. After final round filtering with pae_interaction, relaxed_ddg and sap_score, the sequences were further optimized to have a net-charge of -7.

### Computational design of IGF_EndoTags

To generate flexible IGF-2R agonists, all combinations of Gly-Ser (GS) linkers with various lengths linking D6mb and D11mb were modeled with Alpahfold2. The designs with poor monomer plddt (plddt<85) were dropped.

To generate rigid IGF_EndoTags, the major binding helix from D11mb or the native IGF-2 binding helix, and two interface helices from D6mb were extracted. Both crystal structures obtained for the domain 6 and 11 minibinders in complex with IGF-2 and IGF-2R were used as starting points for design. Domains 6 and 11 of the complex structures were aligned with the respective domains of IGF-2R in the putative receptor internalizing conformation available in the Protein Data Bank (PDB: 6UM2). In this orientation, the two interface helices from the domain 6 binder and the single interface helix from the domain 11 binder were extracted and used as motifs to scaffold by Protein Inpainting^22^. To increase the likelihood of design success, the D11 minibinder structure was adjusted to form an ideal three helical bundle with the two domain 6 helices. Protein Inpainting was implemented such that the interacting residues within 3 angstroms of the receptor maintained the same identity as in the original minibinders. To increase design diversity, the d11 helix motif was randomly perturbed by rigid body translations (up to 5 angstroms) and rotations (up to 10 radians) for each design prior to inpainting a scaffold between the motifs. The best inpainting outputs were selected by RosettaFold LDDT metrics (> 0.5) for the inpainted region and used for sequence design with ProteinMPNN. ProteinMPNN sequence design was performed on the inpainted outputs in their desired complex orientation (with both domains 6 and 11 present) while fixing the original minibinder identities of interface residues (domain 6 minibinder: R4, V8, Q11, D15, V20, K24, M25, I27, I31, and E34, domain 11 minibinder: M1, A4, L7, L8, and W11). After 2000 sequences were generated for each ProteinMPNN input, designs were filtered by predicted Rosetta ddG. AlphaFold 2 structure predictions of the designed sequences were filtered by the pLDDT metric (keeping those with pLDDT > 90), and designs with a subangstrom backbone atom RMSD to the original design models realigned to the D6 and D11 minibinder crystal structures (in complex with the IGFII-M6PR target domains). Finally, the complexes were assessed by Rosetta FastRelax. Designs with ddG metrics less than -40 and Spatial Aggregation Propensity scores less than 35 were selected for expression and experimental assays.

### N-linked glycan verification of epitope

To verify the epitope of the designed binders an N-linked glycan scan was performed. This was performed to rapidly determine if the computational designed binder was interacting with the chosen interface. Four engineered N-linked glycan variants (NN-0975, NN-0979, NN-0981, and NN-0977) with mutation close to the Sortimb binding site were designed and expressed. For design, the computational models were used as a starting point for the computational screen. All positions 10 Å away from the interface were screened using RosettaMatch^43^ followed by a design step to introduce the NXS/T motif into the protein. Computational models were minimized and filtered based on geometrical restraints, CST-score < 5. Next, the four variants were used as bait in the yeast display assay against the computational designed binder, Sortmb, which was displayed on the surface of yeast.

### Yeast surface display screening with FACS

The yeast surface display screening was performed using the protocol as previously described^13,15^. Briefly, DNAs encoding the minbinder sequences were transformed into EBY-100 yeast strain. The yeast cells were grown in CTUG medium and induced in SGCAA medium.

After washing with PBSF (PBS+1% BSA), the cells were incubated with 1uM biotinylated target proteins (IGF-2R, ASGPR, Sortilin) together with streptavidin–phycoerythrin (SAPE, ThermoFisher, 1:100) and anti-c-Myc fluorescein isothiocyanate (FITC, Miltenyi Biotech, 6.8:100) for 30min. After washing twice with PBSF, the yeast cells were then resuspended in PBSF and screened via FACS. Only cells with PE and FITC double-positive signals were sorted for next-round screening. After another round of enrichment, the cells were titrated with biotinylated target protein at different concentrations for 30min, washed, and further stained with both streptavidin–phycoerythrin (SAPE, ThermoFisher) and anti-c-Myc fluorescein isothiocyanate (FITC, Miltenyi Biotech) at 1:100 ratio for 30min. After washing twice with PBSF, the yeast cells at different concentrations were sorted individually via FACS and regrown for 2 days. Next the cells from each subpool were lysated and their sequences were determined next-generation sequencing or Miseq.

For N-linked glycan verification, yeast cells displaying Sortmb were incubated with 100nM N-glycan variants of Sortilin (NN-0975, NN-0979, NN-0981, and NN-0977), separately. Percentage of yeast cells located within the pre-set gate was calculated for each N-glycan variants group and compared with the WT Sortilin group.

### Biolayer interferometry

The binding affinity for the minibinders were determined by the Octet RED96 (ForteBio). To measure the binding affinity, streptavidin-coated biosensors (ForteBio) were first loaded with biotinylated target proteins at 50∼100nM concentration, washed with Octet buffer (10 mM HEPES, 150 mM NaCl, 3 mM EDTA, 0.05% surfactant P20 and 1% BSA), and incubated with titrated concentrations of corresponding binders. To measure the Koff, the biosensors were then dipped back into the Octet buffer. The Kon, Koff and KD were further estimated with the Octet Analysis software.

### Protein and purification

Minibinders and minibinder fusions were expressed in E coli BL21 strain as previously described^13^. Briefly, the DNA fragments encoding the design sequences were assembled into pet-29 vectors via Gibson assembly and further transformed into BL21 strain with heat-shock. Protein expression was induced by the autoinduction system and proteins were purified with Immobilized metal affinity chromatography (IMAC) approach. Next the elutions were purified by FPLC SEC using Superdex 75 10/300 GL column (GE Healthcare). Protein concentrations were determined by NanoDrop (Thermo Scientific) and normalized by extinction coefficients.

Antibody-EndoTag fusions were produced with a mammalian expression system. Light chain of CTX/ATZ antibody and heavy chain fused with EndoTag at C-term constructs were ordered in CMVR from Genscript. Antibody-EndoTag fusions were then expressed via transient co-transfection of the EndoTag-heavy and light chains into Expi293F cells (Life Technologies) via PEI-MAX (Polyscience). In brief, 800mL cultures of Expi293F cells were transfected at a density of 3x10^6 cells per mL of culture using 1 ug of plasmid DNA and 3ug of PEI per mL of culture. These cultures were grown in Expi293F expression medium (Life Technologies) at 37℃ in a humidified, 8% CO2 incubator rotating at 125rpm.

After 6 days of expression, culture supernatants were harvested via 5 minutes of centrifugation at 4000 RCF, 5 minutes of incubation with PDADMAC solution (Sigma Aldrich) added to a final concentration of 0.0375%, followed by an additional 5 minutes of centrifugation at 4000 RCF. Supernatants were clarified via 0.22um vacuum filtration and then treated to a final concentration of 50mM Tris-HCl (pH 8) and 350mM NaCl for IMAC. Gravity IMAC was performed by batch binding the clarified supernatants with 10mL of Ni Sepharose Excel resin (GE Healthcare). After 20-30 minutes of incubation, the resin bed was washed with 10 CV of 20mM Tris-HCl (pH8), 300mM NaCl solution. The proteins were then eluted with 3 CV of 20mM Tris-HCl (pH 8), 300mM NaCl, 300mM imidazole solution. The batch bind process was then repeated with half the amount of resin (5mL) and the eluates from both batch binds were combined. SDS-PAGE was performed on the IMAC eluates to assess purity.

The purified antibody-EndoTag fusions were subsequently concentrated in a 10K MWCO Amicon Ultra centrifugal filter unit (Millipore) and polished via SEC using a Hiload 26/600 Superose 200 column (GE Healthcare) in DPBS (Gibco). The SEC fractions were re-concentrated in the same manner as before to a final concentration of 5 mg/mL. Endotoxin levels were assayed via Endosafe LAL Endotoxin tests (Charles River) and analytical SEC was performed using a Superdex 200 Increase 5/150 column (GE Healthcare) to obtain a high-resolution size profile. Pre- and post-freeze stability was assessed via UV-Vis spectrophotometry as well as SDS-PAGE.

### Cellular uptake evaluation and receptor degradation via flow cytometry

For cellular uptake assays using suspension cell lines (K-562, Jurkat), the cells were incubated with corresponding fluorescence-labeled protein constructs at 37 °C for indicated time, then spun down at 500g for 5min, resuspended and washed with cold PBS. After three washes, the cells were resuspended and transferred to a 96-well plate. For cellular uptake assays using adherent cell lines (U-251MG, Hep3B, Hela, H1975), the cells were incubated with corresponding fluorescence-labeled protein constructs at 37 °C for indicated time, then washed with cold PBS for three times. The cells were then treated with 50uL of trypsin and incubated at 37 °C for 10 min followed by adding 50uL of DMEM media. The resuspended cells were then transferred to a 96-well plate followed by two PBS washes. Flow cytometry was then performed in Attune NxT flow cytometer (Thermo Fisher). The data was analyzed in FlowJo software.

For cell surface receptor degradation experiments, the cells were first incubated with corresponding protein reagents for indicated time at 37 °C, then washed with cold PBS three times. For suspension cell lines, the cells were resuspended and transferred to the 96-well plate; for adherent cell lines, the cells were first treated with trypsin for 10 minutes then transferred to the 96-well plate. The cells were then stained with corresponding fluorescence-labeled antibodies against the corresponding receptor for 1 hour at room temperature. After washing three times with cold PBS for flow cytometry, flow cytometry was performed in Attune NxT flow cytometer (Thermo Fisher). The data was analyzed in FlowJo software.

### Protein degradation via Western blot

Cells were cultured in T75 flasks at 37 °C in a 5% CO_2_ atmosphere. HEP3B (ATCC), HeLa (ATCC), and MDA-MB-231 were cultured in DMEM supplemented with 10% heat-inactivated fetal bovine serum (FBS) and 1% penicillin/streptomycin. Jurkat-CTLA4 and H1975 were cultured in RPMI supplemented with 10% heat-inactivated fetal bovine serum (FBS) and 1% penicillin/streptomycin. Adherent cells were plated (100,000 cells/well in a 24-well plate) one day before the experiment,whereas suspension cells were plated on the day of the treatment. Cells were incubated with 250 µl of complete growth media with pLYTAC or controls for indicated time. Cells were then washed with PBS 3 times and lysed with RIPA buffer supplemented with protease inhibitor cocktail (Roche), 0.1% Benzonase (Millipore-Sigma), and phosphatase inhibitor cocktail (Roche) on ice for 30 minutes. The cells were scraped, transferred to Eppendorf tubes, and spun down at 21,000*g* for 15 minutes at 4 °C. The supernatant was collected and the protein concentration was determined by BCA assay (Pierce). Equal amounts of lysates were loaded onto 4-12% Bis-Tris gel and separated by sodium dodecyl sulfate-polacrylamide gel electrophoresis (SDS-PAGE). Then, the gel was transferred onto a nitrocellulose membrane and stained with REVERT Total Protein Stain (LI-COR), then blocked with Odyssey Blocking Buffer (TBS) (LI-COR) for 1 hour at rt. The membrane was incubated with primary antibodies (rabbit anti-EGFR D38B1 Cell Signaling Technologies, rabbit anti-HER2 2242 Cell Signaling Technologies, rabbit anti-PD-L1 E1L3N Cell Signaling Technologies, rabbit anti-CTAL4 E1V6T Cell Signaling Technologies, mouse anti-vinculin V284 Bio-Rad) overnight at 4 °C, washed 3 times with TBS-T. Subsequently, the membrane was incubated with secondary antibody (800CW goat-anti-mouse or goat-anti-rabbit LI-COR) for 1 hour at rt, and washed 3 times with TBS-T for visualization with an Odyssey CLx Imager (LI-COR). Image Studio (LI-COR) was used to quantify band intensities.

### Fluorescence Imaging

WT HeLa (ATCC CCL-2) were cultured at 37 °C with 5% CO2 in flasks with Dulbecco’s modified Eagle medium (DMEM) (Gibco) supplemented with 1 mM L-glutamine (Gibco), 4.5 g/liter D-glucose (Gibco), 10% fetal bovine serum (FBS) (Hyclone) and 1% penicillin-streptomycin (PenStrep) (Gibco). To passage, cells were dissociated using 0.05% trypsin EDTA (Gibco) and split 1:5 or 1:10 into a new tissue culture (TC)–treated T75 flask (Thermo Scientific ref 156499).

For imaging 35 mm glass bottom dishes were seeded at a density of 20k cells / dish. A final monomeric concentration of 100 nM of ligands were incubated with cultured cells. Cells were fixed 4% paraformaldehyde, permeabilized with 100% methanol, and blocked with PBS + 1% BSA. Cells were immunostained with Anti-LAMP2A antibody (Abcam ab18528) followed by goat anti-rabbit-IgG Alexa Fluor™ 488 secondary antibody (Thermo Fisher A-11034) and 4′,6-diamidino-2-phenylindole (DAPI) (Thermo Fisher D1306) and stored in the dark at 4°C until imaging.

Cells were washed twice with HBSS and subsequently imaged in HBSS in the dark at 37°C. Right before imaging, cells were incubated with 25 µM DTZ. Epifluorescence imaging was conducted on a Yokogawa CSU-X1 microscope equipped with a Hamamatsu ORCA-Fusion scientific CMOS camera and Lumencor Celesta light engine. Objectives used were: 10×, NA 0.45, WD 4.0 mm, 20×, NA 1.4, WD 0.13 mm, and 40×, NA 0.95, WD 0.17–0.25 mm with correction collar for cover glass thickness (0.11 mm to 0.23 mm) (Plan Apochromat Lambda). All epifluorescence experiments were subsequently analyzed using NIS Elements software.

### Confocal microscopy

Indicated cells were seeded in 18 well glass bottom µ-Slides (Ibidi, Cat. No. 81817) at a density of 15k/well. Fluorescencly labeled ligands were incubated with the cultured cells for 0.25, 3, 6 or 24 hours, 30 min before image acquisition, cells were additionally incubated with Lysotracker (ThermoFisherScientific, Cat. No. L7528/L7526/L12492) was added for 30 minutes. Fluorescently labeled anti-IGF-2R (Novus Biological, NB300-514AF647) was added for 30 minutes. Cells were washed 3× in PBS and immediately proceeded to imaging.

Confocal laser scanning microscopy was performed on a Nikon A1R HD25 system equipped with a LU-N4 laser unit (Lasers used: 488 nm, 561 nm, 640 nm). Data was acquired using an 20×, NA 0.75, WD 1.00 mm air objective (Plan Apochromat Lambda) in combination with 1 multialkaline (EM 650 LP) and 2 GaAsP detectors (DM 560 LP EM 524/42 (503-545) and DM 652 EM 600/45 (578-623)). Acquisition was controlled via NIS Elements software and data was analyzed via Fiji and custom-written Python Scripts.

### Mass spectrometry and proteomic

Cell pellets were thawed on ice and lysed in a lysis buffer (400 μL, 1 tablet of Pierce EDTA-free Protease Inhibitor Tablets dissolved in 50 mL of PBS) using a probe sonicator (3 × 3 pulses). Protein concentration was adjusted to 2.0 mg/mL and the samples (100 μL, 200 μg protein) were transferred to new Eppendorf tubes (1.5 mL) containing urea (48 mg/tube, final urea concentration: 8 M). DTT (5 μL, 200 mM fresh stock in H2O, final DTT concentration: 10 mM) was then added to the tubes and the samples were incubated at 65°C for 15 min. Following this incubation, iodoacetamide (5 μL, 400 mM fresh stock in H2O, final IA concentration: 20 mM) was added and the samples were incubated in the dark at 37°C with shaking for 30 min. Ice-cold MeOH (600 μL), CHCl3 (200 μL), and H2O (500 μL) were then added, and the mixture was vortexed and centrifuged (10,000 g, 10 min, 4°C) to afford a protein disc at the interface between CHCl3 and aqueous layers. The top layer was aspirated without perturbing the disk, additional MeOH (600 μL) was added, and the proteins were pelleted (10,000 g, 10 min, 4°C) and used in the next step or stored at −80°C overnight.

The resulting protein pellets were resuspended in EPPS buffer (160 μL, 200 mM, pH 8) using probe sonicator (3 × 3 pulses). Trypsin (10 μL, 0.5 μg/μL in trypsin reconstitute buffer) and CaCl2 (1.8 μL, 100 mM in H2O) were added and the samples were incubated at 37°C with shaking overnight.

Peptide concentration was determined using the microBCA assay (Thermo Scientific) according to the manufacturer’s instructions. For each sample, a volume corresponding to 25 μg of peptides was transferred to a new Eppendorf tube and the total volume was brought up to 35 μL with EPPS buffer (200 mM, pH 8). The samples were diluted with CH3CN (9 μL) and incubated with the corresponding TMT tags (3 μL/channel, 20 μg/μL) at room temperature for 30 min. An additional TMT tag (3 μL/channel, 20 μg/μL, 30 min) was added and the samples were incubated for another 30 min. Labeling was quenched by the addition of hydroxylamine (6 μL, 5% in H2O). Following a 15 min incubation at room temperature, formic acid was added (2.5 μL, final FA concentration: 5%). 20 μL labeled peptides of each channel were combined into a 2.0 mL low-binding Eppendorf tube, and 25 μL of 20% formic acid was added. The resulting mixture was lyophilized to remove the solvents before high pH fractionation.

The spin columns from Pierce High pH Reversed-Phase Peptide Fractionation Kit were pre-equilibrated prior to use. Briefly, the columns were placed in Eppendorf tubes (2 mL), spun down to remove the storage solution (5,000 g, 2 min), and washed with CH3CN (2 × 300 μL, 5,000 g, 2 min) and buffer A (2 × 300 μL, 95% H2O, 5% CH3CN, 0.1% FA, 5,000 g, 2 min). TMT-labeled peptides were re-dissolved in buffer A (300 μL, 95% H2O, 5% CH3CN, 0.1% FA) and loaded onto pre-equilibrated spin columns for high pH fractionation. The columns were spun down (2,000 g, 2 min) and the flow through was used to wash the original Eppendorf tube and passed through the spin column again (2,000 g, 2 min). The column was then washed with buffer A (300 μL, 2,000 g, 2 min) and 10 mM aqueous NH4HCO3 containing 5% CH3CN (300 μL, 2,000 g, 2 min), and the flow through was discarded. The peptides were eluted from the spin column into fresh Eppendorf tubes (2.0 mL) with a series of 10 mM NH4HCO3 / CH3CN buffers (2000 g, 2 min). The following buffers were used for peptide elution (% CH3CN): 7.5, 10, 12.5, 15, 17.5, 20, 22.5, 25, 27.5, 30, 32.5, 35, 37.5, 40, 42.5, 45, 47.5, 50, 52.5, 55, 57.5, 60, 62.5, 65, 67.5, 70, 72.5, 75, 80, 95. Every 10th fraction was combined into a new clean Eppendorf tube (2 mL) and the solvent was removed using a benchtop lyophilizer and stored at -20 °C before analysis.

The resulting 10 combined fractions were re-suspended in buffer A (25 μL) and analyzed on the Orbitrap Fusion mass-spectrometer (4 μL injection volume) coupled to a Thermo Scientific EASY-nLC 1200 LC system and autosampler. The peptides were eluted onto a capillary column (75 μm inner diameter fused silica, packed with C18 and separated at a flow rate of 0.3 μL/min using the following gradient: 5% buffer B in buffer A from 0-10 min, 5%–35% buffer B from 10-129 min, 35%–100% buffer B from 129-130 min, 100% buffer B from 130-139 min, 100%–5% buffer B from 139-140 min, and 5% buffer B from 140-150 min (buffer A: 100% H2O, 0.1% FA; buffer B: 20% H2O, 80% CH3CN, 0.1% FA). Data were acquired using an MS3-based TMT method. Briefly, the scan sequence began with an MS1 master scan (Orbitrap analysis, resolution 120,000, 375−1600 m/z, cycle time 3s) with dynamic exclusion enabled (repeat count 1, duration 30 s). The top precursors were then selected for MS2/MS3 analysis. MS2 analysis consisted of: quadrupole isolation (isolation window was set to 1.2 for charge state z = 2; 0.7 for charge state z = 3; 0.5 for charge states z = 4-6) of precursor ion followed by collision-induced dissociation (CID) in the ion trap (normalized collision energy 35%, maximum injection time 50 ms, MS2 resolution was set to turbo). Following the acquisition of each MS2 spectrum, synchronous precursor selection (SPS) enabled the selection of MS2 fragment ions for MS3 analysis (SPS isolation window was set to 1.3 for charge state z = 2; 0.7 for charge state z = 3; 0.5 for charge states z = 4-6). MS3 precursors were fragmented by HCD and analyzed using the Orbitrap (collision energy 65%, maximum injection time 120 ms). The raw files were converted to mzML files using the MSConvert tool from ProteoWizard (version 3.0.22088). A reverse concatenated, non-redundant variant of the Human UniProt database (2022-11-29) was searched using FragPipe (version 18.0) with the built-in ‘TMT10-MS3’ workflow. The virtual references were used for the data sets due to the lack of a pooled sample. The quantified proteins were filtered with FDR < 1% with median centering normalization. Data are presented as the mean fold change to DMSO-treated controls. n= 3 per group. P values were calculated by a two-tailed unpaired t-test with Welch’s correction.

### IgG/LHDB supernatant clearance assay

Jurkat/K562 cells seeded in 96 culture plates in 300uL media were incubated with AF647 conjugated IgG (Novusbio) or LHDB alone or together with proteinG-EndoTag reagents. At various time points, the cells were pelted down and 30uL of supernatants were extracted and further diluted to 45uL by using a PBS buffer. After shaking in an orbital shaker for 5 minutes, the fluorescence intensity was measured by Neo2 plate reader (BioTek) at wavelength 647 nm. The percentage clearance was measured by normalizing the control group without adding proteinG-EndoTag reagent.

## Acknowledgements

The project or effort depicted was or is sponsored by the Department of the Defense, Defense Threat Reduction Agency grant HDTRA1-21-1-0007 (B.H., L.S.); DARPA Synergistic Discovery and Design (SD2) HR0011835403 contract FA8750-17-C-0219 (W.Y.) and Defense Threat Reduction Agency Grant HDTRA1-21-1-0038 (I.G., W.Y.). This research was supported by the National Institutes of Health’s National Institute on Aging, grant R01AG063845 (B.H., B.C., I.G., W.Y.); the National Institutes of Health’s National Cancer Institute, grant R01CA240339 (I.G.); the Audacious Project at the Institute for Protein Design (L.S., X.W., T.S., I.S., S.W.); the Nordstrom Barrier Institute for Protein Design Directors Fund (B.H., I.G., M.A.); AMGEN Donation to the Institute for Protein Design (S.W., X.W.); The Open Philanthropy Project Improving Protein Design Fund (B.C., I.G.); Dr. Eric and Ms. Wendy Schmidt, and Schmidt Futures funding from Eric and Wendy Schmidt by recommendation of the Schmidt Futures program (I.G.); the European Molecular Biology Organization via ALTF191-2021 (T.S.); NIH grant GM058867 (C.B). The Jane Coffin Childs Memorial Fund for Medical Research (M.Abedi) This research was also funded by National Science Foundation Graduate Research Fellowship and Stanford Center for Molecular Analysis and Design (G.A.). We acknowledge the excellent support from the Biology Imaging Facility at the University of Washington during the confocal imaging experiments. We acknowledge the help from M. Gloegl for SPR experiment setup and M. Exposit for the support of OT-2 liquid handler. We thank X. Li for help with mass spectrometry analysis of proteins; B. Wicky, L. Milles and R. Rogate for optimizing the golden gate assembly protocol. We thank A.Coubet for the analysis of preliminary CryoEM data. We acknowledge D. Lee for the transfection of secretable EGFR-IGF_EndoTag.

## Contribution

B.H., M.Abedi., G.A and B.C. contributed equally. B.H., M.Abedi. and D.B. designed the research. B.H. designed, screened and optimized the binders for IGF-2R and ASGPR. B.C. designed and optimized the binders for Sortilin. I.G. screened and optimized the binders for Sortilin. J.O. and P.G. produced the N-Glycan variants for Sortilin. B.C. and L.Cao. developed the Rifdock binder design pipeline. N.B. set up the PPI MPNN and Alphafold2 pipeline for computational binder design. B.H. and I.S. designed and screened the rigid IGF_EndoTags. B.H. and M.Abedi. studied the endocytosis enhancement effect *in vitro*. B.H., M.Abedi. and G.A. evaluated the protein degradation function of all EndoTags. M.Abedi., J.Z.Z. and T.S. performed the imaging experiments. M.Abedi. evaluated the co-LOCKR AND gate degrader and secretable degrader. R.W. prepared the IGF-2R target protein. R.W. and S.W. performed the CryoEM experiment and collected the CryoEM density map. S.S. wrote the image analysis script used for imaging. Y. W. performed the proteomics analysis for whole-cell protein degradation. B.H., C.W.C, S.C., and S.G produced and purified the protein used in the research. M.Abedi, M.Ahlrichs. and C.D. prepared the cells used for this study. L.Carter. and L.S. coordinated the resources and funding required for the research. C.B. and D.B. supervised this research. B.H., M.Abedi. and D.B. wrote the manuscript with the input from the other authors. All authors analysed data and revised the manuscript.

